# Metabolic flexibility and an unusual route for peptidoglycan muramic acid recycling in mycobacteria

**DOI:** 10.64898/2026.03.23.713626

**Authors:** Stefanos Stravoravdis, Bella R. Carnahan, Rebecca A. Gordon, Kimberly Wodzanowski, Karina Havaleshko, Elisabeth Fils-Aime, Rachel Putnik, Stephen Hyland, Catherine Leimkuhler Grimes, M. Sloan Siegrist

## Abstract

*De novo* biosynthesis of cell wall peptidoglycan is essential for bacterial viability under many growth conditions and is a well-validated antibiotic target. Although generally not essential for bacterial fitness under standard laboratory growth conditions, peptidoglycan recycling can aid bacterial survival under host or antibiotic stress. Peptidoglycan consists of alternating sugars *N*-acetylmuramic acid (Mur*N*Ac) and *N*-acetylglucosamine (Glc*N*Ac) cross-linked by peptides. Recycling of these sugars can proceed via Glc*N*Ac and glucosamine intermediates (*Escherichia coli*-type) or, in the case of Mur*N*Ac, bypass these intermediates altogether (*Pseudomonas*-type). We serendipitously discovered that the pathogen *Mycobacterium tuberculosis* and model organism *M. smegmatis* assimilate 2-modified Mur*N*Ac probes into their peptidoglycan despite lacking the *Pseudomonas*-type machinery that is normally required for incorporation of these molecules. Our data suggest that unmodified and 2-modified Mur*N*Ac incorporate into *M. smegmatis* peptidoglycan via multiple pathways, the former preferentially via an *E. coli*-type route and the latter preferentially via a non-*E. coli*, non-*Pseudomonas*-type route with Glc*N*Ac but not glucosamine intermediates. These findings reveal metabolic flexibility in mycobacterial cell wall recycling that encompasses a previously undescribed pathway.

**Importance:** Cell wall biosynthesis is critical for bacterial replication under many conditions and is a successful drug target. Recycling of cell wall components is often dispensable under laboratory conditions but can promote bacterial survival under stress. We find that mycobacterial species can recycle a cell wall sugar via a mechanism distinct from those used by other bacteria. As pathogenic mycobacteria are subject to stress from the host environment and antibiotics, this mechanism is a potential novel vulnerability for these organisms.

## Introduction

Mycobacteria have a complex cell envelope composed of a cell wall peptidoglycan layer connected via arabinogalactan to an outer mycomembrane (1). Biosynthesis of these layers are successful drug targets e.g., the first-line antituberculars isoniazid (2, 3) and ethambutol (4) disrupt mycomembrane and/or arabinogalactan formation and the second-line drug D-cycloserine disrupts peptidoglycan formation (5, 6). Cell envelope biosynthesis is generally less vulnerable to inhibition when bacteria adapt to stress by slowing replication (7–9). Under these conditions, recycling may become more important. In mycobacteria, for example, recycling of the trehalose sugars in the mycomembrane promote fitness in carbon deprivation (10, 11), hypoxia (12), biofilms (13, 14), host cells (10), and *in vivo* (15). In other bacteria, recycling of the peptidoglycan sugars *N*-acetyl muramic acid (Mur*N*Ac) and *N*-acetylglucosamine (Glc*N*Ac) can support fitness during stationary phase (16); protect against some cell wall-acting antibiotics (17–20); promote biofilm formation (21) and virulence (22–24); and limit immune activation (25). Studies of when, how, and why mycobacteria recycle their peptidoglycan are in their infancy (26–28).

Mur*N*Ac is found only in bacterial peptidoglycan. Two major pathways for recycling this sugar have been described; we refer to them here as *Escherichia coli*- and *Pseudomonas*-type for the organisms in which these mechanisms were discovered. In *E. coli*-type recycling, (anhydro)Mur*N*Ac is released from imported muropeptides and/or imported as a free sugar; phosphorylated sequentially or concomitantly, respectively; converted to Glc*N*Ac and glucosamine intermediates, the latter via removal of the 2-acetyl by NagA (16, 25, 29–37) (**Figure 1**). Glucosamine intermediates can enter central metabolism or be transformed into UDP-Glc*N*Ac and -Mur*N*Ac for *de novo* peptidoglycan biosynthesis (26, 34–36, 38–42). In *Pseudomonas*-type recycling, imported (anhydro)Mur*N*Ac can be transformed more directly into UDP-Mur*N*Ac by AnmK, MupP, AmgK and MurU, bypassing Glc*N*Ac and glucosamine intermediates (17–19, 43–45). Mycobacteria encode homologs for a subset of *E. coli*-type recycling genes, some of which have been experimentally verified, but none for *Pseudomonas*-type recycling genes (26–28). Muramic acid isolated from mycobacterial peptidoglycan can bear either *N*-acetyl or *N*-glycolyl at the 2-position, the proportion of which varies with culture condition (28, 46–48).

**Figure 1.**
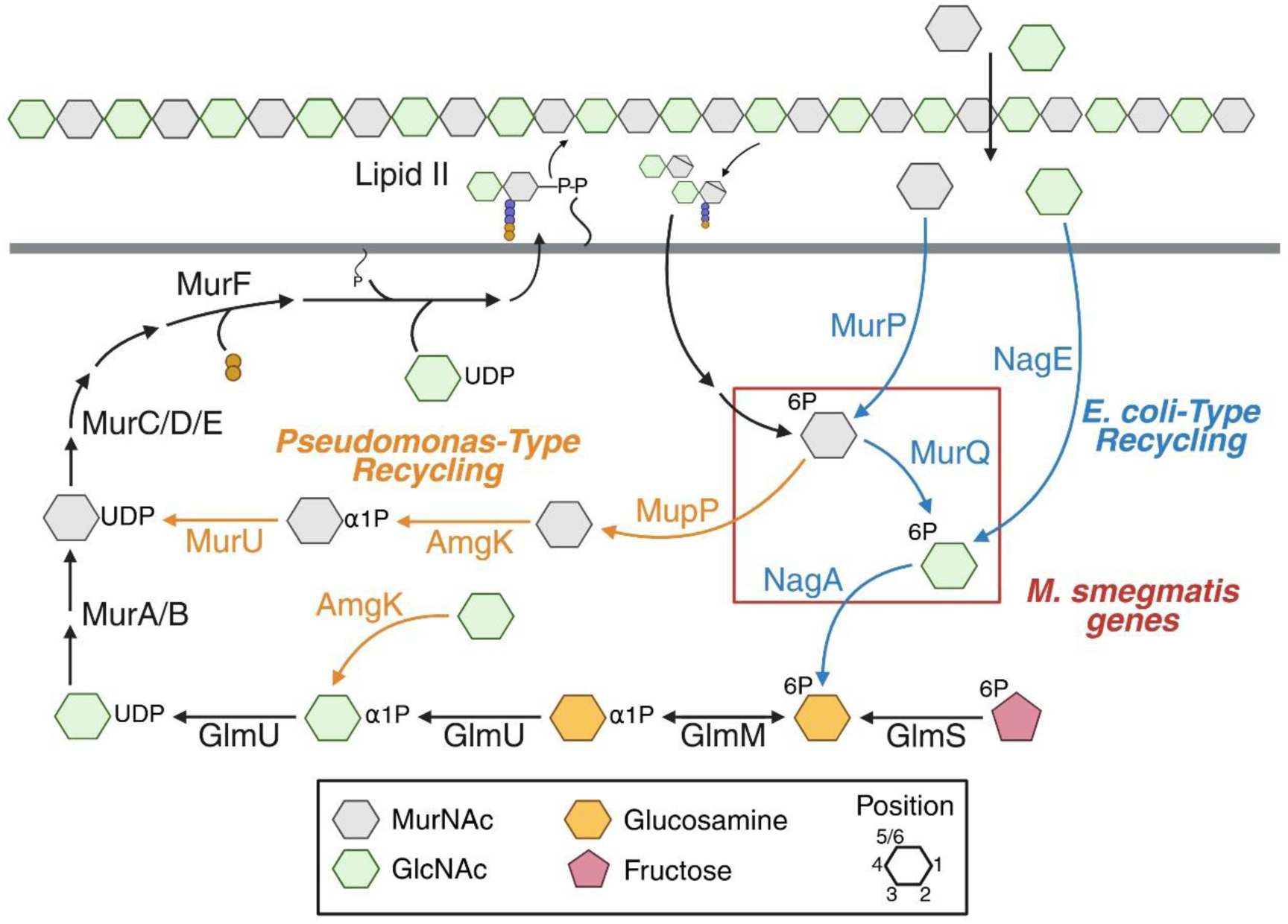
Known routes for Mur*N*Ac recycling and peptidoglycan biosynthesis. In black are steps in *de novo* peptidoglycan biosynthesis. Unique steps of *E. coli*- or *Pseudomonas*-type recycling respectively in blue and orange. *M*. *smegmatis* homologs of *E. coli* recycling pathway indicated with a red box.

Chemical probes have been useful tools for understanding peptidoglycan metabolism and interactions with the host immune system. Metabolic labeling of the carbohydrate core (49) and the stem peptide (50–55) enable downstream detection and manipulation. We previously showed that incorporation of Mur*N*Ac probes functionalized at the 2-position requires endogenous or heterologous expression of *Pseudomonas*-type recycling components AmgK and MurU (50, 51, 55, 56). Unexpectedly, we discovered that we were able to label members of the Mycobacteriales with 2-modified Mur*N*Ac probes despite the absence of *amgK* or *murU* homologs encoded in their genomes and the presence of an *E. coli*-type recycling enzyme NagA, which normally removes the 2-acetyl. We found that 2-alkyne-Mur*N*Ac (2-alkNAM) incorporates into both Glc*N*Ac and Mur*N*Ac isolated from the peptidoglycan of the pathogen *M. tuberculosis* and the model organism *M. smegmatis*. The probe also enhances growth and/or robustly labels the peptidoglycan of *M. smegmatis* defective for *de novo* production of Glc*N*Ac intermediates and, to a lesser extent, glucosamine or Mur*N*Ac intermediates. By contrast, unmodified Mur*N*Ac enhances the growth of *M. smegmatis* defective for *de novo* production of glucosamine and, to a lesser extent, Glc*N*Ac intermediates. Taken together, our data suggest that mycobacteria, in the absence of clear homologs for *Pseudomonas*-type enzymes, can assimilate Mur*N*Ac and related molecules to bypass multiple steps in peptidoglycan synthesis, including steps that *E. coli*- and *Pseudomonas*-type enzymes are not known to bypass.

## Materials and Methods

### Bacterial strains and culture conditions

*M. smegmatis* mc^2^155 (57) and *M. abscessus* ATCC 19977 were grown in Middlebrook 7H9 medium (Fisher Scientific) supplemented with bovine serum albumin (BSA) (2.632% w/v), dextrose (1.053% w/v), sodium chloride (0.447% w/v), glycerol (0.4% v/v), and Tween-80 (0.05% v/v) at 37 °C. *Corynebacterium glutamicum* ATCC 13032 was grown in brain heart infusion (BHI) medium (Becton Dickinson). *M*. *tuberculosis* mc^2^6206 (H37Rv Δ*panCD* Δ*leuCD* (58)) was grown in 7H9 supplemented with 10% Middlebrook Oleic Albumin Dextrose Catalase (OADC), glycerol (0.5% v/v), and Tween-80 (0.05% v/v) at 37 °C. *Escherichia coli lptD*4213, a mutant with an impaired outer membrane (59–62), and *E*. *coli* RFM795 expressing *Tannerella forsythia* AmgK and MurU (55, 56, 59) were grown in lysogeny broth (LB; Genesee Scientific) at 37 °C, while *Arthrobacter globiformis* NRRL B-24224 SEA and *Microbacterium foliorum* NRRL B- 2979 SEA (University of Pittsburgh) were grown in LB at 30 °C.

Libraries of anhydrotetracycline (ATC)-inducible CRISPRi plasmids (built on the pJR962 backbone (63, 64)) and kanamycin- and zeocin-resistant *M*. *smegmatis* deletion mutants were provided by the Mycobacterium Systems Resource Database (MSRdb; (64)). pJR962 empty vector or pJR962 containing d*cas9* and single guide (sg) RNA sequences targeting *msmeg*_1568 (*glmS*), *msmeg*_1559 (*glmM*), *msmeg*_5426 (*glmU*), or *msmeg*_4226 (*murC*) were electroporated into *M. smegmatis* mc^2^155. The sgRNA of *glmU* (gatataatctgggaATGCGCGTGCCGGCTCCGGCGGgtttttgtactcga) was inserted into pJR962 using the NEBridge Golden Gate Assembly Kit (BsmBI-v2) per manufacturer recommendations with a 10:1 ratio of oligo to plasmid. Of note, *M*. *smegmatis* mc^2^155 has two copies of *murA* (*msmeg*_4932), as this gene is found in a duplicated region of the chromosome (65, 66). To enhance our ability to knock down *murA* expression, pJR962 bearing a *murA* sgRNA was electroporated into Δ*murA M*. *smegmatis*, an MSR deletion strain with one copy of *murA.* This generated Δ*murA+*pJR962-sg*murA*, in which one copy of *murA* is deleted and expression of the other can be depleted by CRISPRi.

A knockout of *msmeg*_0193, encoding an ortholog of the peptidoglycan recycling protein MurQ, was generated via the ORBIT strategy detailed by Murphy et al (67). Briefly, wild-type *M*. *smegmatis* mc^2^155 was electroporated with the pKM461 targeting vector. Bacteria were incubated with 25 μg/mL kanamycin and 500 ng/mL ATC until reaching an optical density (OD_600_) of 1.0, after which electrocompetent cells were prepared by washing in increasingly smaller volumes of ice-cold 10% glycerol (Fisher Scientific). Freshly made competent *M. smegmatis* (200 μL) were electroporated with 2 μg of ORBIT oligo (Supplementary Information) and 200 ng of pKM446 plasmid and transferred to 5 mL of 7H9, shaking at 37 °C overnight. The bacteria were centrifuged and plated onto 7H10 agar plates containing 50 μg/mL hygromycin, and successful gene knockout was verified via DreamTaq PCR using manufacturer’s recommendations (Thermo Fisher Scientific; **Table 1**). Colonies of Δ*murQ* were then electroporated with pKM512 for curing of antibiotic resistance. Bacteria were incubated at OD_600_ 0.1 with 500 ng/mL ATC until ∼OD_600_ 1.0, then sub-cultured to repeat this incubation. The bacteria were screened on 7H10 and 7H10 plus hygromycin (50 μg/mL) or zeocin (25 μg/mL) to detect knockout mutants sensitive to all antibiotics. pJR962-sg*glmS* were later introduced into the cured Δ*murQ* strain.

**Table 1.**
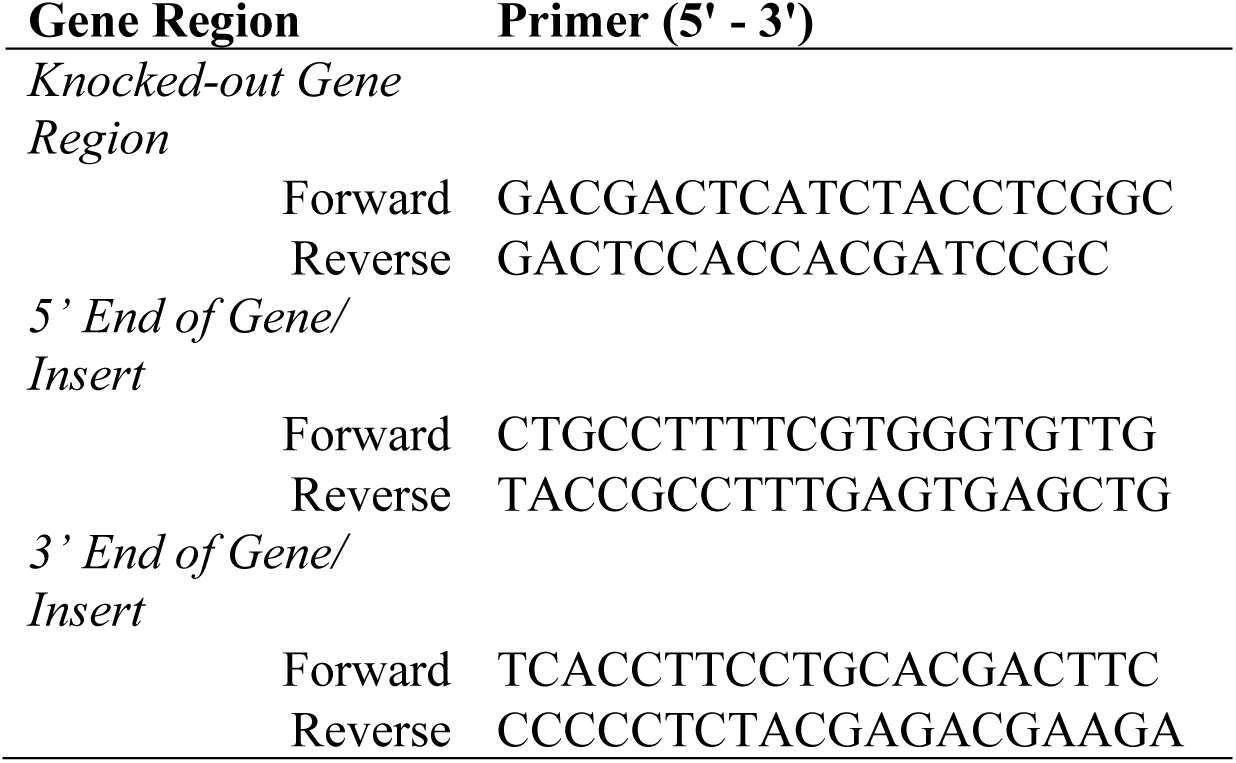
Primers used for identifying MSMEG_0193 knockout Gene Region Primer (5’ - 3’)

### Metabolic labeling for microscopy and flow cytometry

Click reactions were modified from previous protocols (68–71). Bacteria were grown until mid-log phase (OD_600_ ∼0.6-0.8), and treated for 30 minutes with either 2 mM alkyne-D-alanine-D-alanine (alkDADA, also known as EDADA, (69, 72), synthesized by WuXi AppTech), 2-alkyne-or 2-azide-Mur*N*Ac (2-alkNAM or 2-azNAM (50, 51)). Bacteria were centrifuged at 2147 x g for 4 min at 4 °C and washed twice with PBS containing 0.05% Tween 80 (PBST; pre-chilled) using the same conditions. Bacteria were then fixed in 2% paraformaldehyde for 10 minutes, washed thrice with PBST containing 0.01% w/v BSA (PBSTB), and resuspended for copper-catalyzed alkyne-azide cycloaddition (CuAAC). The CuAAC mix consisted of 1 mM copper sulfate, 128 μM Tris(benzyltriazolylmethyl)amine [TBTA], 1.2 mM sodium ascorbate (freshly prepared), and 20 μM fluorescent azide or alkyne (Click Chemistry Tools) in PBSTB diluent. Bacteria were shaken at 600 rpm for 30 minutes using a Benchmark Multi-Therm Heat Shake, then centrifuged, washed twice more with PBSTB and once with PBST, and finally resuspended in PBST prior to imaging on a Nikon Eclipse E600 microscope. Similar reactions were performed in which *M*. *smegmatis* was incubated with 2-tetrazine (Htz)-Mur*N*Ac probe and reacted with trans-cyclooctene (TCO) fluorophore (Click Chemistry Tools) in PBSTB (55). Species other than *M. smegmatis* were grown to mid-log phase (OD_600_ ∼0.6-0.8) and incubated for ∼50% generation time +/- 2 mM alkDADA or 6 mM 2-alkNAM. All subsequent steps were repeated as described above.

Click reactions were performed in duplicate for three biological replicates of the following *M*. *smegmatis* strains: wild-type + pJR962-sg*glmM* or pJR962-sg*glmU* and Δ*murA* + pJR962-sg*murA*. Strains bearing pJR962-sg*glmM* were diluted to OD_600_ ∼0.01 while the remaining strains were diluted to ∼0.02 before either inducing depletion with 50 ng/mL ATC or treating with ethanol (EtOH) as the vehicle control. *Mycobacterium smegmatis* with pJR962-sg*glmM*, pJR962-sg*glmU*, and pJR962-sg*murA* were incubated for 14 hours, 11 hours, or 9 hours, respectively, to induce depletion and trigger a slight decrease in culture turbidity prior to incubation in probe as described above. Median fluorescence intensity was recorded using the 5-laser mode of a BD DUAL LSRFortessa, capturing 30,000 events. The change in fluorescence intensity was computed for each strain relative to the indicated negative controls.

### Phylogenetic tree

A phylogenetic tree was generated from 16S rRNA sequences obtained from NCBI (GenBank: *M. smegmatis*, AJ131761.1; *M. tuberculosis*, MH794239.1; *M. abscessus*, AB548599.1; *C. glutamicum*, AF314192.1; *A. glorbiformis*, AB089841.1; *Microbacterium foliorum*, AJ249780.1; *E. coli*, AB681728.1) using the List of Prokaryotic names with Standing in Nomenclature as guidance (73). A neighbor-joining tree was constructed with 1000 bootstrap replicates via MEGA12 using default settings.

### Mass spectrometry

A 10 mL culture of wild-type *M*. *smegmatis* was grown to OD_600_ ∼0.6, diluted four-fold, then split into 10 mL aliquots, each incubated shaking for 3 hours at 37 °C +/- 2 mM of alkyne-D-alanine (alkDA; R-propargylglycine; Thermo Fisher, Waltham, MA) or 5 mM 2-alkNAM. Bacteria were centrifuged at 3000 x g for 5 min, washed twice in 10 mL of PBS, then resuspended in 2.5 mL of digestion buffer (250 µL of 1 M Tris pH 7.9, 25 μL of 5 M NaCl, 20 μL of 0.5 M EDTA, and 4.75 mL of deionized water). Freshly prepared lysozyme (Sigma-Aldrich; 50 mg/mL, 20 μL) was added to each sample. Bacteria were shaken at 37 °C for 3 days with additional infusions of 20 μL of freshly prepared lysozyme every 12 hours. To collect the peptidoglycan fragments released by lysozyme, bacteria were centrifuged at 3075 x g for 5 seconds and added to an Amicon Ultra 3K Filter. Samples were centrifuged at 3000 x g for 30 min to collect flowthrough. The filter was washed with 2 mL of deionized water and spun as described prior to collect the flowthrough. All flowthroughs were combined, flash frozen on dry ice, and lyophilized overnight. A similar protocol was performed for *M*. *tuberculosis*, except that bacteria were incubated with probes for 3 days (∼3.5 doublings). The lyophilized samples were dissolved in 50 µL of deionized water and loaded onto Acquity UPLC BEH C18 column 2.1 x 50 mm (Waters) using a S11 Dionex UHPLC. The samples were eluted with the following gradient: 0.5 mL/min – 1 linear gradient starting from 0% eluent A (0.1% formic acid in water) to 50% eluent B (formic acid in acetonitrile) in 6 min and subsequently underwent high-resolution mass analysis (when needed desired fragments were subjected to MS-2) on a Q-Exactive Orbitrap (Thermo Fisher Scientific). All data were processed and analyzed on a Thermo Xcalibur Qual Browser.

### MurNAc, GlcNAc, and 2-alkNAM rescue experiments

Primary cultures of CRISPRi plasmid-bearing *M. smegmatis* were grown in 7H9 medium supplemented with 25 µg/mL kanamycin (and 25 µg/mL zeocin for Δ*murA+*pJR962-sg*murA*). Bacteria were back-diluted and grown overnight in fresh 7H9 medium without antibiotic to mid-log phase (OD_600_ ∼0.4-0.8). After adjusting the OD_600_ to 0.05 in 7H9 (0.005 for pJR962-sg*glmM*), *M. smegmatis* was grown in triplicate in 96-well flat-bottom plates +/- 50 ng/mL ATC to induce expression of d*cas9* and sgRNAs. Plates were continuously shaken at 37 ℃ in a Biotek Synergy 2 plate reader at medium setting with hourly OD_600_ readings for 20 hours. At 0 and 4 hours, 5 mM Mur*N*Ac or Glc*N*Ac (Sigma-Aldrich) or vehicle control was added to the appropriate wells. Mur*N*Ac was prepared according to literature precedent and matched reported spectrum (74) or purchased from Sigma-Aldrich. We observed that elevated concentrations of Mur*N*Ac (> 5 mM) dampen growth of wild-type *M. smegmatis* and *M. tuberculosis*, a phenotype that has been reported previously in other species (**Figures S1A-S1B**; (16, 75)). Thus, we opted to use 5 mM Mur*N*Ac for most *M. smegmatis* rescue experiments. Additional growth curves were generated for pJR962-sg*glmS*, pJR962-sg*glmM*, pJR962-sg*glmU*, and Δ*murA* + pJR962-sg*murA*, treating the bacteria at 0 and 4 hours with 0 mM or 5 mM Mur*N*Ac, Glc*N*Ac, or 2-alkNAM. Growth was monitored using the Biotek Synergy 2 plate reader.

### Quantitative real-time PCR (qRT-PCR)

qRT-PCR was performed similarly to Prithviraj et al (76). Wild-type *M. smegmatis* containing the pJR962 empty vector, pJR962-sg*glmM*, or pJR962-sg*glmU* were grown in 100 mL cultures +/- 50 ng/mL of ATC until OD_600_ ∼0.4-0.6. Twenty-five OD units were pelleted for each treatment at 3200 x g for 10 min. The supernatant was discarded, and the pellets were resuspended in 1 mL of TRIzol (Thermo Fisher Scientific) and transferred to tubes containing 0.5 mL of zirconia beads (Benchmark Scientific). Bacteria were bead-beat for 1 min with 2 min intervals on ice before incubating at room temperature for 5 min. The samples were spun for 30 seconds at 10,000 x g. The top TRIzol layer was transferred to a fresh RNAse-free tube, treated with 200 µL chloroform, shaken vigorously for 15 seconds, and left at room temperature for 10 min. The samples were spun at 12,000 x g for 10 min at room temperature and the top aqueous layer was saved. To each tube, 0.1 volumes of 3 M sodium acetate (pH 5.2, made with DEPC water) and 0.7 volumes of isopropanol were added. The tubes were inverted and stored at −20 °C for 2 hrs to precipitate nucleic acids. The samples were centrifuged at 16,000 x g for 30 min at 4°C. The pellets were saved and washed twice in ice cold 70% ethanol (DEPC) using the same spin conditions. Samples were dried and the nucleic acids were resuspended in 40 µL DEPC-water (Sigma-Aldrich), 4 µL 10X TURBO DNase I buffer (Thermo Fisher Scientific), and 1 µL 1 U/µL DNase I (Thermo Fisher Scientific). The samples were vortexed and briefly spun at maximum speed before incubating for 2 hrs at 37 °C. RNA was further concentrated using Zymo RNA Clean and Concentrator following the manufacturer’s instructions (Zymo Research) and stored at −80 °C until cDNA synthesis. To generate cDNA, the different samples were processed using Invitrogen’s SuperScript IV Reverse Transcriptase according to manufacturer’s instructions, save that 50 µM random hexamers (New England Biotech), 1 µL of Promega RNasin Plus RNA inhibitor (Fisher Scientific), and 2 µM gene-specific reverse primers for *glmM*, *glmU*, and *gyrB* (housekeeping control (77)) were added (**Table 2**).

**Table 2.**
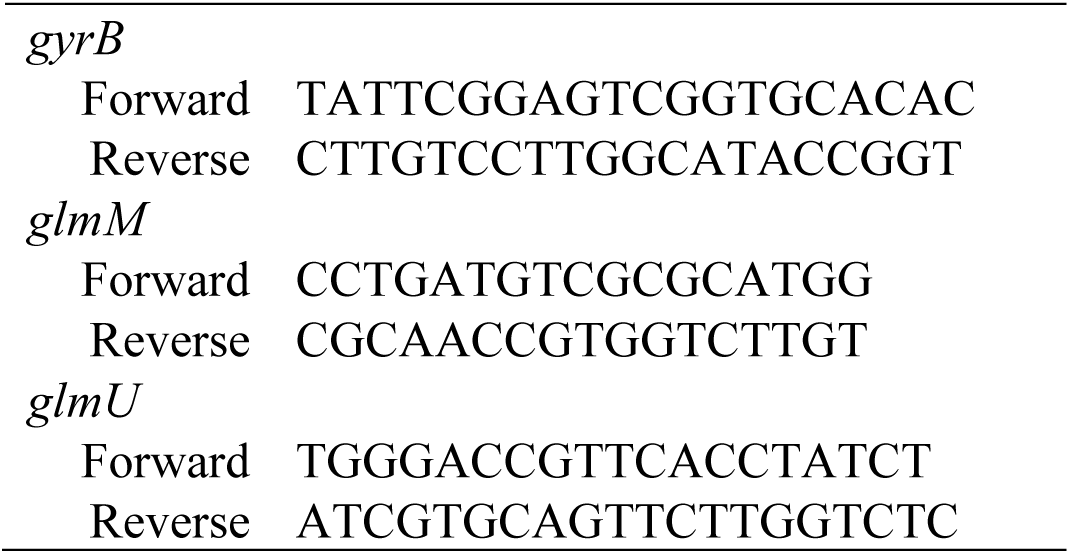
Primers used for RT-qPCR reaction Gene Primer (5’ - 3’)

Each 20-µL reaction was composed of 10 µL ITaq Universal SYBR Green Supermix (Bio-Rad) with cDNA (final concentration 50 ng/µL) and appropriate primers (final concentration 200 nM; **Table 2**). Quantitative RT-PCR was performed on a DNA Engine Opticon 2 from MJ Research (Saint-Bruno-de-Montarville, Quebec). Reaction conditions were performed in technical triplicate as follows: 95 °C for 5 min; 40 cycles of incubations at 95 °C for 30 s, 67 °C for 30 s, taking reads after each cycle. A melting curve was performed from 50 °C to 95 °C. The 2^−ΔΔCT^ was calculated using C_T_ as the threshold cycle.

## Results

### Corynebacterineae cell surface labeling via 2-modified MurNAc probes

Classic (anhydro) Mur*N*Ac recycling, first defined in *E. coli* (35), proceeds via (anh)Mur*N*Ac → Mur*N*Ac-6-P → Glc*N*Ac-6-P then loss of the *N*-acetyl at the 2-position, mediated by NagA, to furnish glucosamine-6-P (Glc*N*-6-P; **Figure 1**). Glc*N*-6-P can then be funneled into glycolysis or into peptidoglycan biosynthesis, the latter occurring via Glc*N*-6-P → Glc*N*-1-P → Glc*N*Ac-1-P → UDP-Glc*N*Ac → UDP-Mur*N*Ac (**Figure 1**). An alternative Mur*N*Ac recycling pathway (18, 19, 24, 43, 78), first defined in *Pseudomonas* species, bypasses both Glc*N*Ac and Glc*N* intermediates and proceeds via Mur*N*Ac-6-P → Mur*N*Ac → Mur*N*Ac-1-P → UDP-Mur*N*Ac, the latter two steps mediated by AmgK and MurU (**Figure 1**). In both the *E. coli*- and *Pseudomonas*-type systems, the nucleotide sugars are processed by additional enzymes to produce the final, lipid-linked precursor lipid II.

We previously synthesized Mur*N*Ac probes functionalized at the 2-acetyl with azide, alkyne, or minimal tetrazine (50–53, 55) and showed that peptidoglycan labeling of bacteria is dependent on the endogenous or heterologous expression of AmgK and MurU. Presumably NagA either does not act on the probes, or deacetylation and potentially other metabolic transformations render the chemical handles undetectable. Surprisingly, we were able to label the model organism *Mycobacterium smegmatis* using these 2-modified Mur*N*Ac probes (50–53, 55) followed by detection via copper-catalyzed alkyne-azide cycloaddition (CuAAC) or tetrazine ligation to fluorescent labels (**Figures 2, S2**). This was an unexpected phenotype because mycobacterial species lack obvious homologs for AmgK and MurU (27). By contrast, and as expected, *E. coli* labeling occurred only upon heterologous expression of *amgK* and *murU* (**Figure 2**). Mur*N*Ac probe labeling of *M. smegmatis* was spatially like that reported for other cell envelope probes (11, 54, 69–71, 79–89), including an alkyne derivative of D-alanine-D-alanine (52, 72) (alkDADA; **Figure 2**). More specifically, fluorescence derived from 2-alkNAM or alkDADA was enriched at the septa and poles, respectively the locations of cell division and elongation in mycobacteria (69, 81, 90–95). These data suggest that the Mur*N*Ac probes label in the vicinity of active *M. smegmatis* cell envelope metabolism via an unknown mechanism.

**Figure 2.**
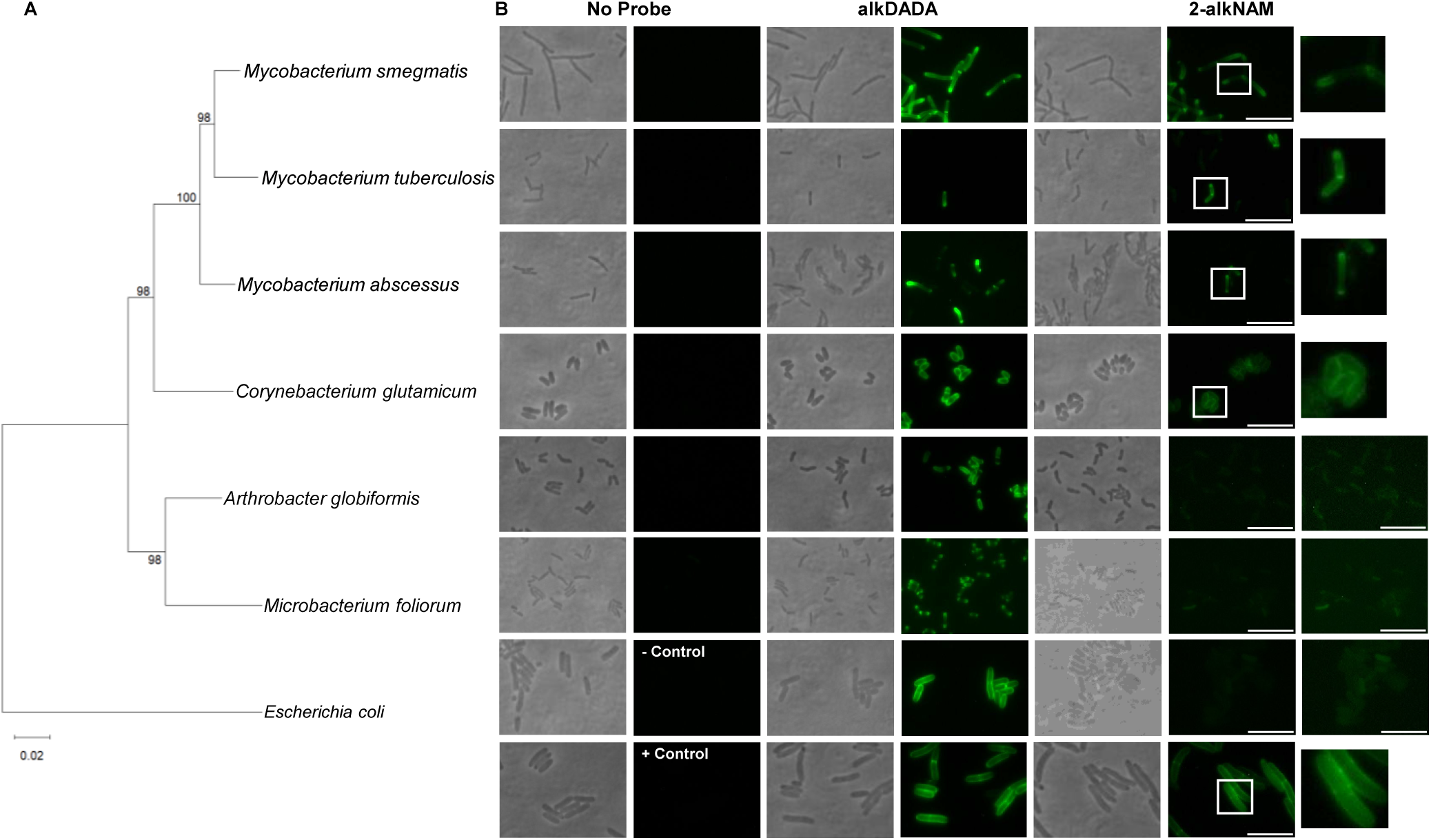
Cell surface labeling phenotypes across the *Actinomycetes* with cell wall probes alkDADA (69, 72, 111) or 2-alkNAM (50, 51). (A) Phylogenetic tree of 16S rRNA sequences. (B) Representative images of bacteria incubated or not with 2 mM alkDADA or 6 mM 2-alkNAM probes (∼25% [*M. smegmatis*] or ∼50% generation time [all other species]) and subjected to CuAAC with fluorescent azide label. The negative control for 2-alkNAM labeling is *E. coli*, a species that lacks *amgK and murU* homologs; we used the *lptD*4213 mutant, a strain with a known outer membrane defect (59–62), to enhance permeability of the label (61) and increase sensitivity of detection. The positive control for 2-alkNAM labeling is *E. coli* strain RFM795 (55, 56, 59) which heterologously expresses AmgK and MurU from *Tannerella forsythia*. Zoomed in images of labeled cells or high-exposure FITC images were shown for Mur*N*Ac-treated bacteria (*far right*). Scale bars, 8 μm.

We next investigated the phylogenetic distribution of Mur*N*Ac probe labeling in species related to *M. smegmatis* that also lack AmgK and MurU homologs. Members of the Mycobacteriaceae family were incubated with alkDADA or 2-alkNAM prior to CuAAC (**Figure 2**).

*Mycobacterium tuberculosis* and *M*. *abscessus* labeled with both probes at the poles and septa, like *M*. *smegmatis*, although there remained sizable subpopulations of unlabeled bacteria, particularly for *M*. *abscessus* (**Figure 2**). We previously reported cell-to-cell labeling heterogeneity in *M. tuberculosis* for other cell envelope probes that require a fluorophore-appending step for detection (62, 69); we speculate that the phenotype is due to fluorophore permeability rather than underlying cell envelope metabolism. Mur*N*Ac probes also labeled *Corynebacterium glutamicum*, albeit with non-polar localization that was distinct from that of other chemical probes for peptidoglycan (68, 82, 96–99) (**Figure 2**). As *C. glutamicum* is a member of the Corynebacteriaceae family but in the same Mycobacteriales order as the mycobacterial species, we next tested whether the Mur*N*Ac labeling phenotype extended to a different order, the Micrococcales. *Arthrobacter globiformis* and *Microbacterium foliorum* labeled at their poles and septa with alkDADA but had only background fluorescence when treated with 2-alkNAM (**Figure 2**). These data suggest that Mur*N*Ac probes can label different members of the Mycobacteriales order and that labeling is spatially coincident with sites of active cell envelope metabolism in the Mycobacteriaceae family.

## 2-modified MurNAc probe incorporates into mycobacterial peptidoglycan MurNAc and GlcNAc

Mycobacterial species have a homolog of *E. coli*-type NagA but not *Pseudomonas*-type AmgK nor MurU (26–28). Therefore, these organisms lack a clear mechanism for Mur*N*Ac probe assimilation that would preserve chemical handles installed at the 2-position. However, the septal and polar localization of labeling in *M. smegmatis, M. tuberculosis,* and *M. abscessus* suggested that Mur*N*Ac probes incorporate near or at sites of active cell envelope metabolism. We therefore tested directly whether 2-alkNAM incorporates into mycobacterial peptidoglycan. We treated *M. smegmatis* and *M. tuberculosis* with 2-alkNAM, subjected the bacteria to lysozyme, an enzyme that hydrolyzes β-(1,4) linkages between Mur*N*Ac and Glc*N*Ac, and analyzed by LC-MS. We found that the alkyne was present in lysosome-digested peptidoglycan fragments for both organisms (**Figures 3A-3B, S3-S8**). Unexpectedly, some of the digested fragments contained both alkyne and *N*-glycolyl modifications, suggesting that the Glc*N*Ac was modified with the alkyne while the Mur*N*Ac retained the canonical *N*-glycolyl modification (**Figures 3Ai, 3Bii-3Biii, S3, S7-S8**). Thus, *M. smegmatis* and *M. tuberculosis* can assimilate 2-alkNAM into both Mur*N*Ac and Glc*N*Ac of their peptidoglycan.

**Figure 3.**
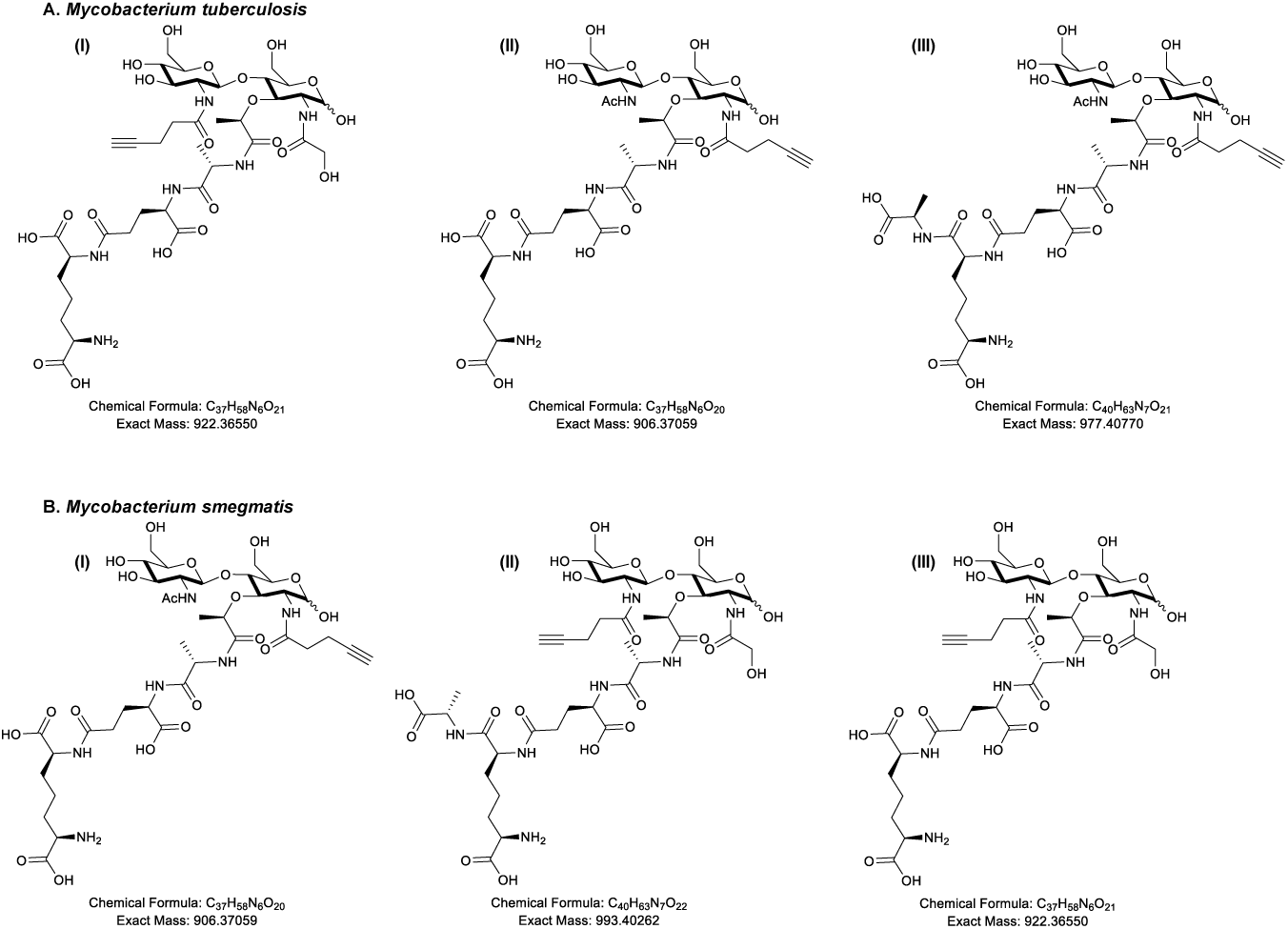
Disaccharide Fragment found from purified and digested *M. tuberculosis* (A) and *M. smegmatis* sacculi (B). Structures and exact masses are shown from LC-MS in *M. tuberculosis* and *M. smegmatis* for muropeptides released after incubation in 2-alkNAM. Presence of the alkyne handle was detected in *N*-glycolyl-modified disaccharide in Glc*N*Ac (3Ai, 3Bii-3Biii) as well as in Mur*N*Ac (3Aii-Aiii, 3Bi) for both species. Complete spectra in Supplementary Information.

### E. coli-type recycling assimilates MurNAc better than 2-modified MurNAc probe in M. smegmatis

The presence of both alkyne-labeled Mur*N*Ac and Glc*N*Ac in the peptidoglycan of 2-alkNAM-labeled *M*. *smegmatis* and *M. tuberculosis* (**Figure 3**) implied that recycling of 2-modified Mur*N*Ac into peptidoglycan can proceed via Glc*N*Ac intermediates, as UDP-Glc*N*Ac is converted into UDP-Mur*N*Ac and both are then used to construct lipid II. However, this was a puzzling inference as mycobacteria encode *E. coli*-type NagA, which removes the 2-acetyl of Mur*N*Ac to generate Glc*N*-6-P en route to UDP-Glc*N*Ac, and *M. smegmatis* additionally encodes *E. coli*-type MurQ, which converts Mur*N*Ac-6-P to Glc*N*Ac-6-P to generate the substrate for NagA. We therefore hypothesized 2-modified Mur*N*Ac recycling in mycobacteria partially or completely avoids glucosamine intermediates, thus enabling the metabolite to retain a chemical handle. We reasoned that *E. coli*-type recycling may assimilate Mur*N*Ac poorly in the absence of *murQ* (only *M. smegmatis*) and/or when the 2-position is modified (all mycobacterial species). Because *E*. *coli*-type recycling of exogenous Mur*N*Ac generates Glc*N*-6-P, it can bypass the requirement for *de novo* synthesis of this metabolite by the essential enzyme GlmS (**Figure 1**, (16, 26, 34–36, 40–42)). We therefore used CRISPRi to knock down *glmS* in wild-type or Δ*murQ M. smegmatis* and asked whether we could rescue bacterial growth by addition of Mur*N*Ac or 2-alkNAM to the growth medium. In this system, the expression of both dCas9 and the guide RNA is under the control of an anhydrotetracycline (ATC)-inducible promoter (63, 64, 100). Depletion of peptidoglycan biosynthetic enzymes results in low turbidity readings with high variability, the latter of which we attribute to debris from lysed cells, e.g., **Figures 4** and **5**. Consistent with a functional *E. coli*-type recycling pathway, we found that exogenous Mur*N*Ac rescues growth of *M. smegmatis* carrying a *glmS*-targeting guide in the presence of ATC (**Figure 4A**). Rescue was lost in the absence of *murQ* (**Figure 4A**) and diminished when the 2-position of Mur*N*Ac was modified with an alkyne (**Figure 4C**). These data suggest MurQ-dependent, *E. coli*-type recycling is operative in *M. smegmatis* and assimilates Mur*N*Ac with an unmodified 2-acetyl better than it does 2-modified MurNAc.

**Figure 4.**
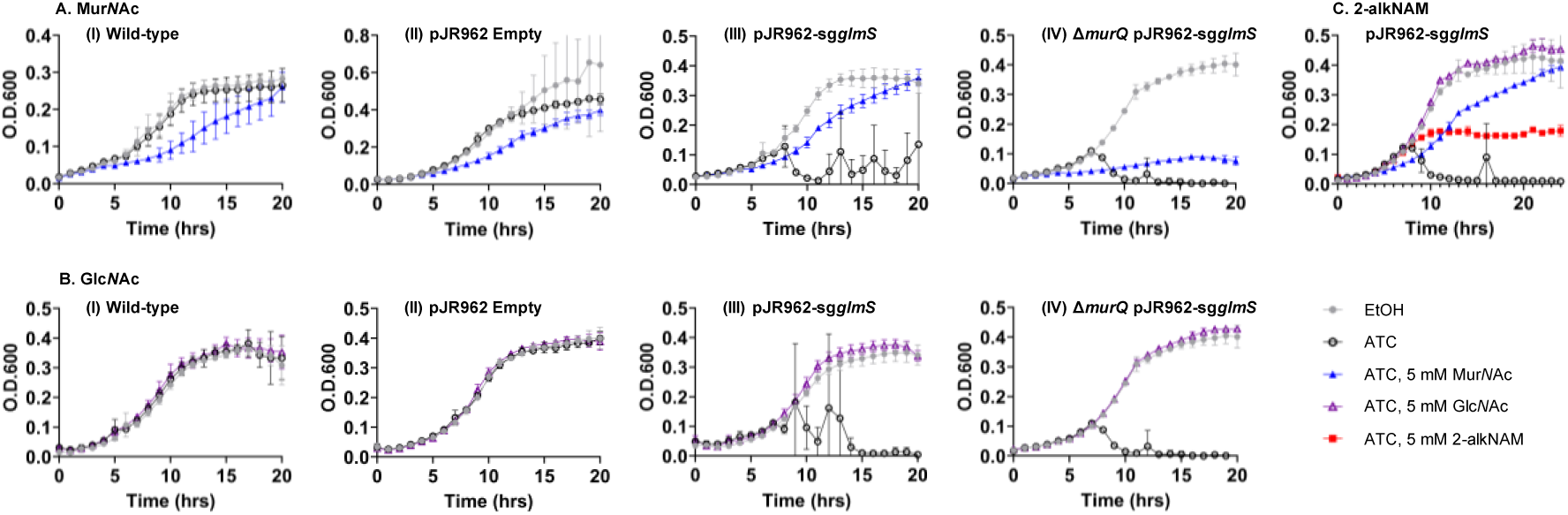
*E. coli*-type recycling assimilates Mur*N*Ac, Glc*N*Ac, and, to a lesser extent, 2-modified Mur*N*Ac probe. Wild-type or Δ*murQ M. smegmatis* +/- pJR962 plasmid (empty) or containing *glmS* guide (sg*glmS*) and *dcas9* under an ATC-inducible promoter were treated +/-50 ng/mL ATC or ethanol (EtOH) carrier control and supplemented with the indicated amounts of Mur*N*Ac (A) or Glc*N*Ac (B) at 0 and 4 hours of growth. Representative data shown from two biological replicates performed in technical triplicate. *M. smegmatis* pJR962-sg*glmS* was additionally treated +/- 50 ng/mL ATC or EtOH and supplemented with Mur*N*Ac, Glc*N*Ac, or 2-alkNAM (C) as described above. Representative data shown from two biological replicates performed in technical duplicate.

**Figure 5.**
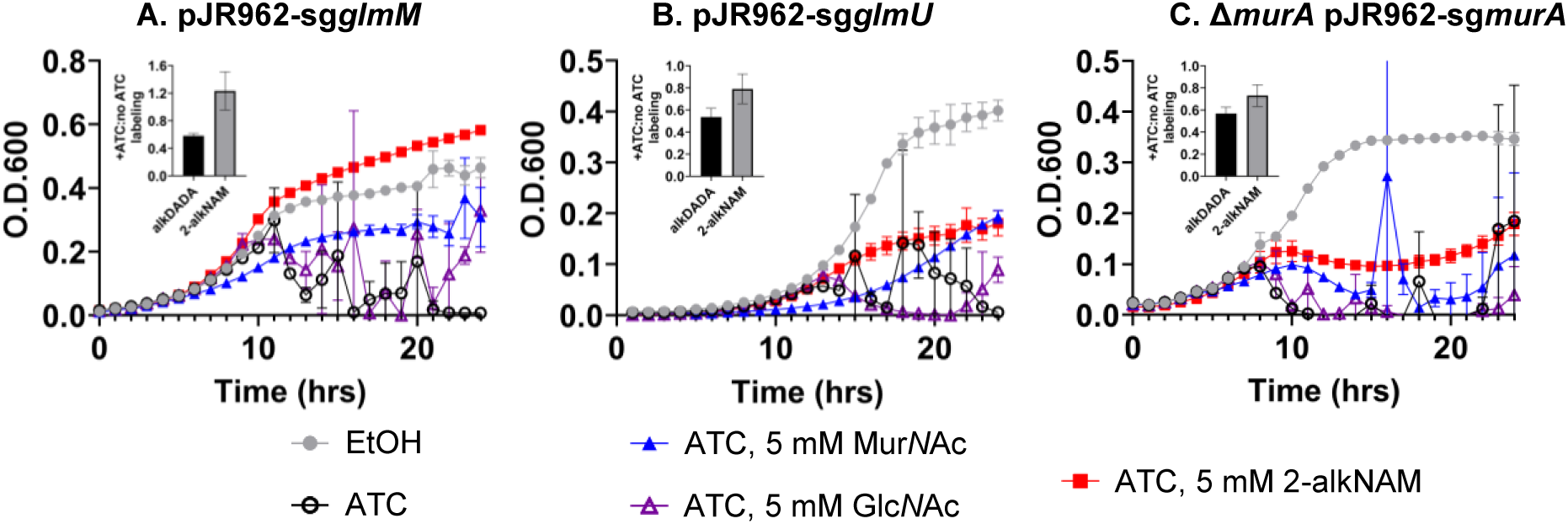
2-modified Mur*N*Ac probe is primarily assimilated into *M. smegmatis* peptidoglycan by non-*E. coli*, non-*Pseudomonas*-type recycling pathway. Wild-type or Δ*murA M. smegmatis* +/- pJR962 plasmid containing *dcas9* and guides for *glmM* (sg*glmM*; (A)), *glmU* (sg*glmU*; (B)), or *murA* (sg*murA*; (C)) guides under an ATC-inducible promoter were treated +/- 50 ng/mL ATC or ethanol (EtOH) carrier control and supplemented with 5 mM of Mur*N*Ac, Glc*N*Ac, or 2-alkNAM at 0 and 4 hours of growth. Representative data shown from two biological replicates performed in technical duplicate or triplicate. Insets show metabolic labeling with alkDADA and 2-alkNAM probes at 1 hour after first evidence of drop in bacterial turbidity. Presence of probes were detected by CuAAC ligation to azide-fluorophore after fixation. Ratios of median fluorescence intensities +/- ATC plotted from representative of at least two biological replicates performed in technical triplicate.

### E. coli-type recycling assimilates GlcNAc in M. smegmatis

*M. smegmatis*, in contrast to other mycobacterial species, can recycle Glc*N*Ac and encodes the proteins to do so via an *E. coli*-type pathway that does not require MurQ: Glc*N*Ac → Glc*N*Ac-6-P (NagE) → Glc*N*-6-P (NagA; **Figure 1** (26–28)). Consistent with an operative pathway for *E. coli*-type Glc*N*Ac recycling, we found that exogenous Glc*N*Ac could rescue growth of *glmS*-depleted *M. smegmatis* but, in contrast to Mur*N*Ac, did so in a MurQ-independent fashion (**Figure 4B**).

### Non-E. coli, non-Pseudomonas-type recycling pathway assimilates 2-modified MurNAc probe better than MurNAc in M. smegmatis

While our growth rescue experiments suggested that 2-alkNAM could transform via *E. coli*-type recycling into Glc*N*-6-P (**Figure 4C**), our analysis of peptidoglycan from 2-alkNAM-treated *M. smegmatis* and *M. tuberculosis* was consistent with a pathway that uses Glc*N*Ac but not glucosamine intermediates (**Figure 3**). We therefore hypothesized that 2-modified Mur*N*Ac could also access the *de novo* pathway for peptidoglycan synthesis at one or more steps after GlmS production of Glc*N*-6-P but prior to MurA production of UDP-Mur*N*Ac (**Figure 1**). Glc*N*-6-P is converted to Glc*N*-α-1P by GlmM then to Glc*N*Ac-α-1P and finally UDP-Glc*N*Ac by the bifunctional enzyme GlmU (101, 102)). Bypass of GlmM or GlmU depletion by exogenous Mur*N*Ac would respectively support models for recycling of the sugar into Glc*N*Ac-α-1P or UDP-Glc*N*Ac. Using CRISPRi to knock down GlmM or GlmU in *M. smegmatis* (**Figure S9A-S9B**), we found that addition of exogenous 2-alkNAM partially or completely rescues ATC-mediated GlmM or GlmU depletion, respectively (**Figure 5A-5B**). Together with our biochemical data (**Figure 3**) these data suggest that 2-modified Mur*N*Ac can be recycled into Glc*N*Ac intermediates.

In contrast to 2-alkNAM, unmodified Mur*N*Ac and Glc*N*Ac completely bypass GlmS (**Figure 4**), suggesting that they primarily transform into Glc*N*Ac-6-P followed by Glc*N*-6-P via *E. coli*-type recycling pathways. Consistent with this interpretation, exogenous Mur*N*Ac rescued growth of GlmM- or GlmU-depleted *M. smegmatis* to a lesser extent than 2-modified Mur*N*Ac, and exogenous Glc*N*Ac did not rescue at all (**Figure 5A-5B**).

We considered the possibility that our growth rescue experiments might be confounded by metabolic dysfunction. Specifically, we observed that elevated concentrations of Mur*N*Ac (> 5 mM) dampen growth of wild-type *M. smegmatis* and *M. tuberculosis*, a phenotype that has been reported previously in other species (**Figures S1A-S1B**; (16, 75)) and may be due to sugar phosphate toxicity (103–110). As an orthogonal test of whether 2-alkNAM could bypass GlmM or GlmU depletion, we asked whether the probe was able to incorporate into peptidoglycan using cell surface labeling as a readout. We and others have shown that dipeptide D-amino acid probes incorporate into peptidoglycan and/or lipid II. Specifically, incorporation of dipeptide probes occur via the last cytoplasmic step of the lipid II biosynthetic pathway, MurF (69, 72, 111, 112). We found that 2-alkNAM labeling was dampened less than alkDADA labeling upon depletion of GlmM or GlmU (**Figure 5A-5B**). The persistence of peptidoglycan labeling and enhanced growth in GlmM- or GlmU-depleted *M. smegmatis* supplemented with 2-alkNAM are consistent with the presence of the alkyne in Glc*N*Ac and Mur*N*Ac in mycobacterial peptidoglycan (**Figure 3, S3-S8**). Together these data suggest that the probe is recycled directly into Glc*N*Ac intermediates.

### Pseudomonas-like recycling pathway partially assimilates 2-modified MurNAc probe in M. smegmatis

After GlmM and GlmU generate UDP-Glc*N*Ac, MurA converts the nucleotide sugar to UDP-Mur*N*Ac. In *Pseudomonas*-type recycling, Mur*N*Ac can be directly assimilated into UDP-Mur*N*Ac and therefore bypass MurA and avoid Glc*N*Ac and Glc*N* intermediates ((18, 19, 24, 43, 78) and **Figure 1**). We had assumed, given the lack of obvious *Pseudomonas*-type enzymes encoded by mycobacterial species, that MurNAc and 2-modified MurNAc probe would be unable to bypass MurA. Contrary to this assumption, however, we found that 2-alkNAM could partially rescue the growth of MurA-depleted *M. smegmatis* (**Figure 5C**). As well, 2-alkNAM labeling was dampened less than labeling by alkDADA under this condition (**Figure 5C**).

## Discussion

Previous *in silico* analysis and experimental characterization (26–28) revealed that mycobacteria encode a limited subset of *E. coli*-type recycling proteins for peptidoglycan sugars, e.g., NagA, NagC, NagZ/LpqI, and, in the case of *M. smegmatis*, MurQ. Beyond these studies, the mechanisms by which mycobacteria recycle Mur*N*Ac are largely unclear. Given the presence of NagA, which deacetylates the 2-position of Mur*N*Ac, and the apparent absence of a *Pseudomonas*-type recycling pathway to bypass MurA/MurB (lack of AmgK or MurU homologs and **Figures 5**), we were initially surprised that various mycobacterial species incorporate 2-modified Mur*N*Ac probes into their peptidoglycan (**Figures 2, 3, S2**). The presence of the alkyne handle from 2-alkNAM in both Mur*N*Ac and Glc*N*Ac (**Figures 3, S3-S8**) and the ability of 2-alkNAM and unlabeled Mur*N*Ac to respectively rescue *glmU* fully or partially (**Figure 5B-5C**) together suggest that recycled Mur*N*Ac can enter *de novo M. smegmatis* peptidoglycan biosynthesis as UDP-Glc*N*Ac (**Figure 6**).

**Figure 6.**
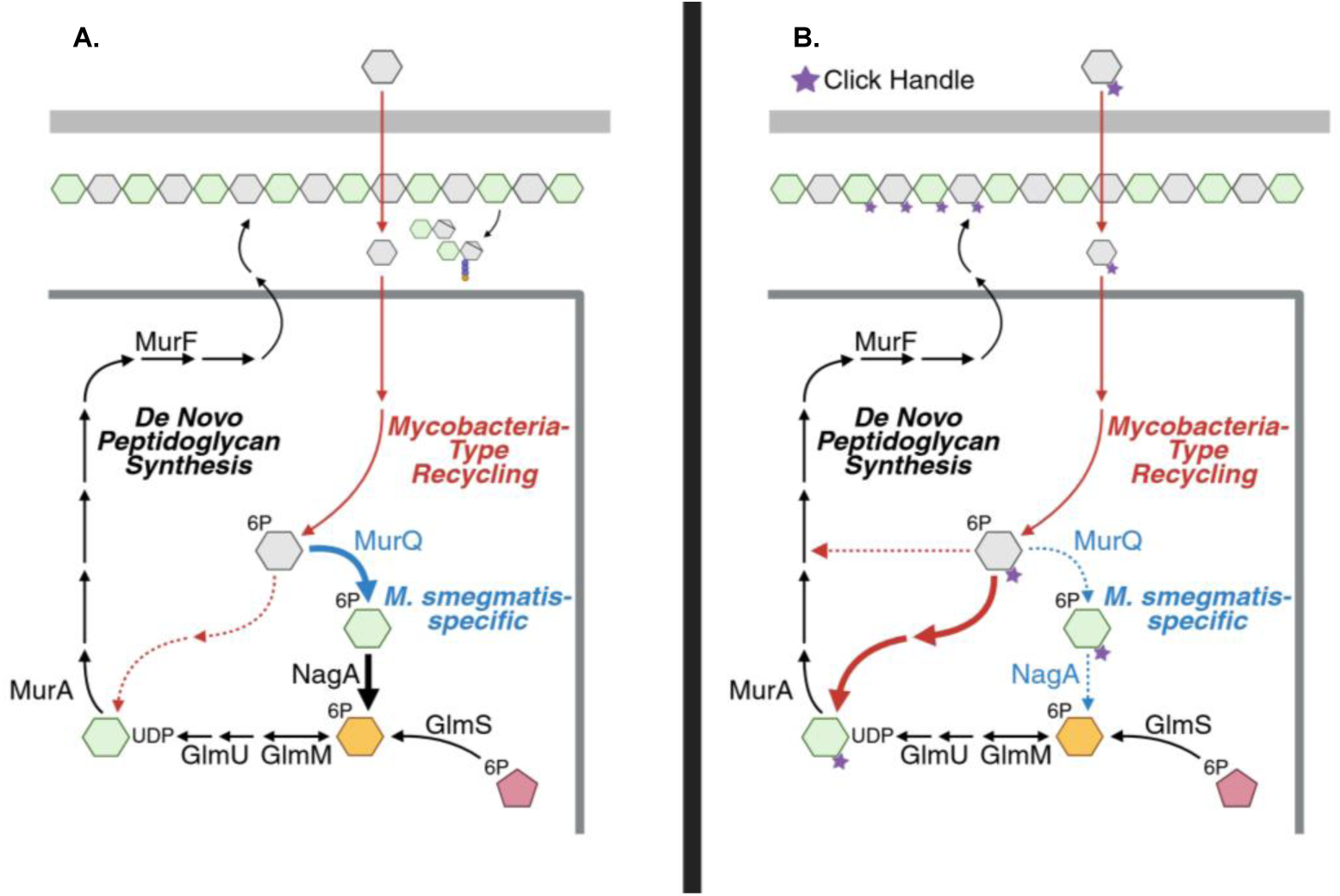
Multiple pathways for Mur*N*Ac recycling in *M. smegmatis*. (A) Assimilation of unmodified Mur*N*Ac. (B) Assimilation of 2-modified Mur*N*Ac. Mycobacterial species other than *M. smegmatis* lack MurQ but can incorporate 2-modified Mur*N*Ac probe into their peptidoglycan (Figure 3) and/or cell surfaces (Figure 2) suggesting conservation of the pathway in (B). Thicker arrows indicate preferred paths by which Mur*N*Ac or 2-modified Mur*N*Ac are funneled back into *de novo* synthesis.

*M*. *smegmatis* has additional *E. coli*-type recycling proteins that are not present in other members of the Mycobacteriales, including NagE, AnmK, MurK, MurQ, and NagB (27, 43). However, *M. tuberculosis* and *M. abscessus* also label with Mur*N*Ac probes and do so at the known polar and septal sites of cell envelope metabolism (**Figure 2**). For *M. tuberculosis* we have confirmed that the alkyne handle from 2-alkNAM is present in peptidoglycan fragments. Thus, we favor a model in which Mur*N*Ac probes are incorporated into peptidoglycan via a similar mechanism across mycobacteria.

*C. glutamicum* also labels with Mur*N*Ac probes, but in contrast to other cell envelope probes, including D-amino acid dipeptides (69, 72, 111), the fluorescence is evenly distributed around the cell periphery (**Figure 2**). We previously reported that peptidoglycan probes localize in part to the sidewall in mycobacterial species, in addition to the poles and septa, and that sidewall labeling is enhanced upon cell wall damage (69–71). We speculate that the early steps of Mur*N*Ac recycling in *C. glutamicum* are like those of mycobacterial species but that the later steps are divergent, such that Mur*N*Ac probe labeling in *C. glutamicum* reports cell wall repair instead of, or in addition to, cell wall expansion. It is also possible that labeling is non-specific in this organism. Given that we have not yet tested these hypotheses, we refer to the Mur*N*Ac recycling pathway(s) described herein as “Mycobacteria-type” instead of “Mycobacteriales-type.”

Mur*N*Ac can be used as a sole carbon source in different mycobacterial species, including *M. tuberculosis* and *M. smegmatis* (27). Mur*N*Ac processing likely occurs via a mechanism in which an unidentified lactyl etherase acts on the sugar to produce Glc*N*Ac and lactate, the latter entering central metabolism. The fate of the released Glc*N*Ac and site of its release (cytoplasm vs. periplasm) are not yet known. In contrast to *M. smegmatis*, the other Mycobacteriales tested—*M. tuberculosis* and *M. abscessus*—cannot use Glc*N*Ac as a sole carbon source, cannot assimilate radiolabeled Glc*N*Ac, and/or do not encode homologs of the inner membrane Glc*N*Ac transporter/kinase NagE (27, 28, 113). Nonetheless, all three species label with 2-modified Mur*N*Ac probes (**Figures 2, 3**, **S2**) and the peptidoglycan of 2-alkNAM-treated *M. smegmatis* and *M. tuberculosis* contains the alkyne handle in Glc*N*Ac in addition to Mur*N*Ac. Thus, we speculate that a common pathway for funneling recycled Mur*N*Ac back into peptidoglycan involves uptake of Mur*N*Ac across the inner membrane by an unknown transporter followed by lactyl etherase generation of Glc*N*Ac and/or its intermediates in the cytoplasm.

In *E. coli*-type recycling, Mur*N*Ac is converted into both Glc*NA*c and glucosamine intermediates prior to peptidoglycan incorporation: MurP transports and phosphorylates Mur*N*Ac to Mur*N*Ac-6-P, MurQ removes the lactate to Glc*N*Ac-6-P, NagA deacetylates to Glc*N*-6-P, and GlmM and GlmU convert to UDP-Glc*N*Ac (**Figures 1, 6A-6B**) (16, 26, 33–42). Recycled Glc*N*Ac is also converted into glucosamine intermediates: NagE transports and phosphorylates the sugar to Glc*N*Ac-6-P, followed by NagA, GlmM, and GlmU conversion to UDP-Glc*N*Ac (29–32, 36). We find that *glmS* depletion is rescued by either Mur*N*Ac (unmodified, and to a lesser extent, the 2-modified probe) or Glc*N*Ac (**Figure 4**) but that *glmM* and *glmU* depletion are rescued only by Mur*N*Ac (the 2-modified probe, and to a lesser extent, unmodified; **Figure 5A-5B**). We speculate that in *M. smegmatis*, Glc*N*Ac is recycled as it is in *E. coli*, i.e. transported and phosphorylated by NagE to Glc*N*Ac-6-P. However, none of the Mycobacteriales tested here encode MurP homologs, and only *M. smegmatis* encodes MurQ. We therefore posit that a common pathway for at least a portion of recycled Mur*N*Ac to enter back into peptidoglycan synthesis includes transport and transformation of the sugar to a Glc*N*Ac intermediate other than Glc*N*Ac-6-P via a lactyl etherase other than MurQ, followed by additional, unidentified enzyme(s) that generate UDP-Glc*N*Ac (**Figure 6**).

Muramic acid isolated from mycobacterial peptidoglycan can bear either *N*-acetyl (Mur*N*Ac) or *N*-glycolyl (Mur*N*Glyc; (28, 46–48)). Our data suggest that in *M. smegmatis* Mur*N*Ac and 2-modified probe preferentially access different pathways for incorporation, with the former primarily assimilated by an *E. coli*-type pathway with glucosamine intermediates and the latter primarily assimilated by a non-*E. coli*, non-*Pseudomonas* type pathway with Glc*N*Ac intermediates. We speculate that mycobacteria may have evolved multiple pathways to accomodate differential recycling of Mur*N*Ac and Mur*N*Glyc. Given that peptidoglycan glucosamine is known to be 2-modified with *N*-acetyl (Glc*N*Ac) but not *N*-glycolyl (Glc*N*Glyc), it is possible that *N*-glycolyl on recycled Mur*N*Glyc may be replaced by *N*-acetyl during the non-*E. coli*, non-*Pseudomonas* recycling process. Additionally, or alternatively, the partial rescue of *murA*-depleted *M. smegmatis* by 2-modified probe (**Figure 5C**) suggests that mycobacteria may harness a third potential recycling route (**Figure 6B**) that could enable Mur*N*Glyc to avoid both glucosamine and Glc*N*Ac intermediates, potentially enabling the assimilation directly into UDP-Mur*N*Glyc.

Cell envelope salvage and recycling can be vulnerable pathways for stressed, non/slow-replicating bacteria. In *M. tuberculosis*, for example, recycling of trehalose sugars from the outer mycomembrane occurs in *in vitro* models of stress, including hypoxia (12), biofilms (13, 14), and carbon deprivation (10, 11), and promotes survival in host macrophages (10), survival *in vivo* (15), and drug tolerance (114). Less is known about when and why mycobacteria recycle peptidoglycan sugars (26–28, 115). In other species, cell wall recycling mutants often do not have phenotypes under standard laboratory conditions (116, 117) but can have defects in stationary phase (16); sensitivity to certain cell wall-acting antibiotics (17–20); impaired biofilm formation (21); attenuation in vivo (22, 23) and in macrophages (24); abnormal immune activation (25). Our data suggest that mycobacteria can harness a Mur*N*Ac recycling pathway that is distinct from both of the previously-characterized *E. coli*- and *Pseudomonas*-type mechanisms. Future identification of the transporters and enzymes that mediate the Mycobacteria-type pathway prior to its intersection with *de novo* peptidoglycan biosynthesis will enable dissection of physiological significance during stress and evaluation as a potential therapeutic target.

## Acknowledgements

We thank the directors of the University of Massachusetts Amherst Flow Cytometry and Biophysical Characterization core facilities, Drs. Amy Burnside and Stephen Eyles, for equipment use and advice, as well as the Massachusetts Life Science Center for the funding for the equipment, and the lab of Dr. Klaus Nusslein for use of their DNA Engine Opticon 2. These studies were supported by the National Institutes of Health: U01 CA221230 to C.L.G. and M.S.S.; R21 AI163949 to M.S.S. and C.L.G; National Research Service Award T32 GM139789 to S.S.; 9R01AI191445; 5T32GM133395; 3P20GM104316; and R21 AI188455 to M.S.S.

## Conflict of Interest

M.S.S. is a co-founder and acting CSO of Latde Diagnostics.

